# Improving the quality of combined EEG-TMS neural recordings: Introducing the Coil Spacer

**DOI:** 10.1101/170431

**Authors:** K.L. Ruddy, D.G. Woolley, D. Mantini, J.H. Balsters, N. Enz, N. Wenderoth

## Abstract

**Background:** In the last decade, interest in combined transcranial magnetic stimulation (TMS) and electroencephalography (EEG) approaches has grown substantially. Aside from the obvious artifacts induced by the magnetic pulses themselves, separate and more sinister signal disturbances arise as a result of contact between the TMS coil and EEG electrodes.

**New method:** Here we profile the characteristics of these artifacts and introduce a simple device – the coil spacer - to provide a platform allowing physical separation between the coil and electrodes during stimulation.

**Results:** EEG data revealed high amplitude signal disturbances when the TMS coil was in direct contact with the EEG electrodes, well within the physiological range of viable EEG signals. The largest artifacts were located in the Delta and Theta frequency range, and standard data cleanup using independent components analysis (ICA) was ineffective due to the artifact’s similarity to real brain oscillations.

**Comparison with Existing Method:** While the current best practice is to use a large coil holding apparatus to fixate the coil ‘hovering’ over the head with an air gap, the spacer provides a simpler solution that ensures this distance is kept constant throughout testing.

**Conclusions:** The results strongly suggest that data collected from combined TMS-EEG studies with the coil in direct contact with the EEG cap are polluted with low frequency artifacts that are indiscernible from physiological brain signals. The coil spacer provides a cheap and simple solution to this problem and is recommended for use in future simultaneous TMS-EEG recordings.

## INTRODUCTION

There has been a recent surge in the number of publications reporting simultaneous transcranial magnetic stimulation (TMS) and electroencephalography (EEG) recordings. This amalgamation of methods has introduced valuable new ways to probe and measure the brain, such as with TMS evoked responses (eg. Ferreri et al. 2011, Miniussi and Thut 2010, Bonato et al. 2006) and TMS induced oscillations (eg. Paus et al. 2001). While the vast majority of studies have focused on post-TMS EEG signals, emerging theories on state-based stimulation make the claim that differences in ongoing neural oscillations at the moment when brain stimulation occurs likely impact outcome measures (see Thut and Pascual-Leone 2010). For these investigations, the EEG signal measured *before* the TMS pulse contains critical information.

Considering the immense methodological challenges posed by the application of high intensity magnetic pulses during the recording of delicate low amplitude EEG signals, it is not surprising that the focus of most attempts to improve combined TMS-EEG protocols has been on the substantial signal disturbances caused in the immediate interval following the pulse. In this regard, much progress has been made (Veniero et al. 2009,Virtanen et al. 1999, Mutanen et al. 2013, Rogasch et al. 2013, Julkunen et al. 2008), but there remains another less obvious source of artifact that has received little attention and is crucial to the study of pre-TMS brain states. This is the signal disturbance that arises simply from contact between the TMS apparatus and the surface of the EEG cap. In the absence of a dedicated investigation comparing signals with and without this disturbance, the extent of the artifact and its impact upon resulting interpretations of data remains unknown. Movement artifacts are in the frequency range of bioelectric events, making them particularly difficult to discern from true brain signals, posing a high risk of polluting the EEG in a way that is disguised as viable physiological data. Here we focus specifically on the artifact associated with direct contact between the TMS coil and electrodes during simultaneous EEG recording, and introduce a simple solution to improve the quality of such recordings for future investigations.

## METHODS

### Participants

Six healthy volunteers (age: 22-29, 4 male, 3 female) participated in the study. All gave informed consent to procedures. The experiments were approved by the Kantonale Ethikkommission Zürich, and were conducted in accordance with the Declaration of Helsinki (1964).

### Experimental setup and procedure

Subjects sat in a comfortable chair with both arms and legs resting in a neutral position supported by foam pillows. Surface electromyography (EMG, Trigno Wireless; Delsys) was recorded from right First Dorsal Interosseous (FDI) and Abductor Digiti Minimi (ADM). EMG data were sampled at 2000 Hz (National Instruments, Austin, Texas), amplified and stored on a PC for off-line analysis.

### Combined TMS-EEG

TMS was performed with a figure-of-eight coil (internal coil diameter 50 mm) connected to a Magstim 200 stimulator (Magstim, Whitland, UK). Prior to application of the EEG cap, the ‘hotspot’ of the right FDI was determined as the location with the largest and most consistent MEPs, and was marked directly onto the scalp with a skin marker. The TMS coil was hand held over this location with the optimal orientation for evoking a descending volley in the corticospinal tract (approximately 45 degrees from the sagittal plane in order to induce posterior-anterior current flow). Once the hotspot was established, the EEG cap (Electrical Geodesics Inc. (EGI), Oregon, USA) was applied and electrodes were filled with gel. Through the EEG cap, the previously marked position of the FDI hotspot was located visually and the TMS coil was applied directly over this point. With the coil directly resting on the EEG cap, the lowest stimulation intensity at which MEPs with a peak-to-peak amplitude of approximately 50 μV were evoked in at least 5 of 10 consecutive trials was taken as Resting Motor Threshold (RMT). The procedure to establish RMT was repeated again with the introduction of the coil spacer between the cap and the TMS coil.

### The Coil Spacer

The coil spacer (Figure 1A & B) is a plastic circular tripod (1.1 cm in height) with a 12.5 cm handle, which was 3D printed (Ultimaker 2, design files available online at https://3dprint.nih.gov/discover/3dpx-007789) and can be customized to virtually every EEG cap and TMS coil available. The three conical feet attached to the circular ring are wider at the bottom than the top, to spread pressure widely over the scalp area and avoid discomfort. The circular ring is hollow in the middle to allow direct vision for positioning the centre of the ring on top of the marked hotspot. To ensure accurate placement of the coil over the hotspot, a red line is marked on the spacer handle, which should be aligned with the middle of the top rim of the TMS coil, in order to ensure that the centre of the coil (at the intersection of the two electromagnetic coils, where the magnetic pulse is strongest) is placed directly over the hotspot (which is positioned in the centre of the spacer ring).

**Figure 1.**
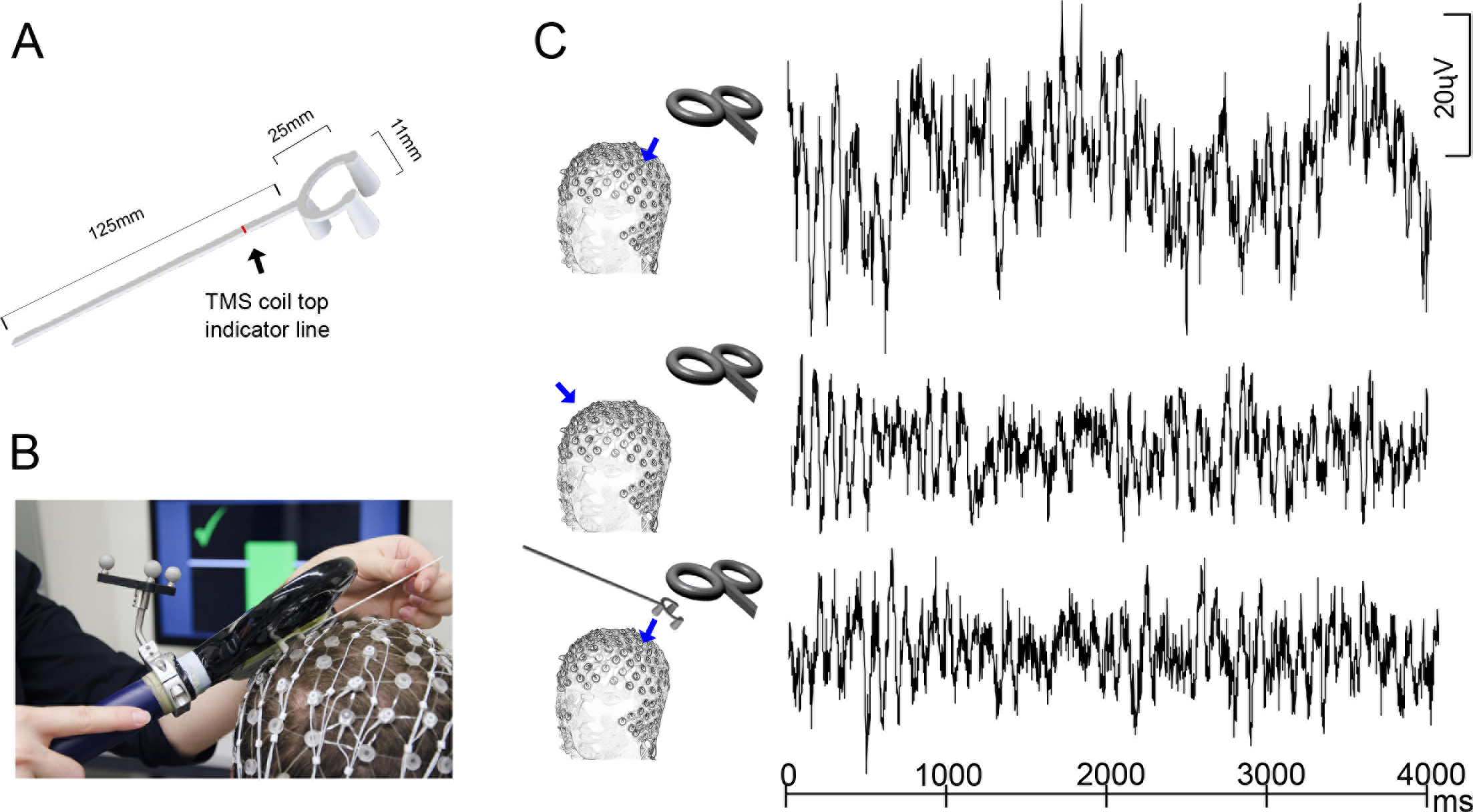
Combined TMS-EEG using the Spacer. The spacer design (A) and an image of the spacer in use during an experiment (B). EEG recordings from a representative subject of one 4000 ms epoch in each condition (C): Blue arrows indicate the location of the electrode from which recordings are displayed. In the upper panel, the TMS coil is placed directly on top of the EEG cap over the left hemisphere motor hotspot during recording. The mid panel depicts the same epoch of data but recorded from the corresponding electrode on the right hemisphere while the TMS coil is on the left hemisphere. The lower Panel shows the left hemisphere hotspot recording when the spacer is placed between the TMS coil and the EEG cap.

EEG signals were recorded inside an electromagnetically shielded room, with a 64 channel gel-based TMS-compatible cap (MicroCel, EGI). The TMS unit was positioned outside the room, with the coil cable passed inside via a wave guide. Signals were amplified and sampled at 1000 Hz. The channel closest to the TMS hotspot was noted for later analyses. Impedances were monitored throughout and maintained below 50 kΩ.

There were two blocks of simultaneous TMS and EEG recording, each containing 20 magnetic pulses at an intensity of 120% RMT, with a 6-8 second inter-stimulus interval between pulses (jittered to avoid anticipation effects). In one block, TMS was applied while the coil was in direct contact with the EEG cap on the head, and in the other block the spacer was placed between the coil and the head in order to provide a platform over the electrodes, meaning that the coil could ‘hover’ over the cap without directly touching electrodes. Resting EEG data without TMS was also collected.

### EEG data processing, analysis and statistics

As the focus of this investigation was upon the quality of EEG signals recorded while the TMS coil is placed on the head but *before* the TMS pulse was applied, the first step was to extract 4000 ms epochs of data in the interval immediately before the magnetic pulse. By epoching in this way, the large artifacts associated with the pulses are excluded from any further analysis, and normal filtering procedures can be applied to the remaining data. Thus, our investigation focused on artifacts that are associated with direct contact between the TMS coil and the EEG electrodes.

After epoching, the same EEG data were analysed in two different ways using EEGLAB (Delorme and Makeig 2004). First, signals were analysed in their ‘raw’ state, with only minimal processing applied (a 0.1-80Hz bandpass filter, and average re-referencing). Then, separately the signals were processed and cleaned using Independent Components analysis, and bad components containing physiological artifacts were identified by correlations with signals recorded from electrooculographic (EOG) and facial EMG electrodes. The rejection of artifactual bad components using this method was automated and therefore not prone to subjective experimenter bias, and consistent across datasets. Bad channels were detected and interpolated. The purpose of this dual approach displaying raw and cleaned data was to demonstrate explicitly the profile of the TMS-EEG movement artifacts in their unaltered form, and subsequently demonstrate whether traditional processing approaches are capable of rendering the data useable.

Power spectral density was computed for both raw and cleaned signals for the data recorded from the electrode closest to the TMS hotspot, and for a ‘control’ electrode, which was selected as the corresponding location in the opposite hemisphere. This electrode was chosen as it could be expected to demonstrate similar amplitude signals to the hotspot electrode in the hemisphere where TMS was applied, but will be minimally affected by the application of TMS on the opposite side of the head. In order to justify this choice of control electrode, an additional analysis was conducted to compare power spectra from the two chosen locations to verify that no differences exist at rest (Supplementary Table 1).

EEG signals from the electrode closest to the TMS ‘hotspot’ were compared to signals from the corresponding electrode in the opposite hemisphere. A power spectrum was computed and decomposed into the following frequency bands: delta (1-4 Hz), theta (5-7 Hz), alpha (8-14 Hz), beta (15-30 Hz) and gamma (31-80 Hz). Log transformed average power values within each band were entered into a repeated measures ANOVA model, with a 2×2 full factorial design. The factors were ‘electrode’ (two levels: hotspot or control), and ‘presence of spacer’ (two levels: ‘spacer’ or ‘no spacer’). ANOVA models were conducted for each of the five frequency bands, and separately for raw and cleaned data. Partial eta squared (*η*^2^) effect sizes are reported for main effects, where greater than 0.14 is considered a large effect.

### TMS unit noise test

In order to further isolate the source of the artifact arising from contact of the TMS coil on the EEG cap, we conducted a separate study with one subject resting with eyes open. We collected continuous EEG data, while turning the TMS unit on and off at 20 second intervals (randomized ON and OFF conditions, controlled using custom MATLAB software and an Arduino interface to the TMS unit). This test was repeated in separate blocks using the Spacer and with No Spacer. For each block we separated this data into 40 4-second epochs (in order to use identical EEG processing pipeline as for the main experiments), 20 of which occurred with the TMS unit switched ON and the remaining 20 with it switched OFF (randomized ON OFF conditions). We conducted power spectral analyses on this data comparing TMS unit ON and OFF conditions for Spacer and No Spacer in order to demonstrate whether there are differences in the power spectrum when the TMS coil is touching the head that may simply arise from electrical noise being conducted through the TMS equipment.

## RESULTS

With the addition of the spacer, resting motor thresholds increased on average by 11±1% of the maximum stimulator output (MSO), compared to using no spacer (without spacer mean 47±9 MSO, with spacer mean 58±8 MSO).

Upon inspection of the data it is clear (Figure 1C & 2A) that the amplitude of signals collected without the spacer in the delta frequency range are an order of magnitude greater compared to those collected with the inclusion of the spacer, or compared to the control electrode in the opposite hemisphere. The movement artifacts manifest in the power spectrum predominantly in the lowest frequency ranges. At the lowest frequencies that we were capable of extracting given the 4000ms epochs, (0.25-1Hz) power values were on average 10.3 times greater when no spacer was used compared to when the spacer was in place preventing the coil from contacting the cap. At 1Hz the signals were 8.3 times greater. In the delta range these signals were 6.5 times larger on average, theta 1.5 times, alpha 1.5 times, beta 1.1 times and gamma 1.2 times larger.

For EEG signals in the delta frequency range, there was a significant ‘electrode’ by ‘spacer/no spacer’ interaction (Figure 2B), indicating that the amplitude of 1-4Hz oscillations were significantly higher when the TMS coil was in contact with the EEG cap compared to when the spacer was used, and this difference was only present at the hotspot electrode and not at the control electrode. Importantly, this interaction was present both in the raw data (F[1,5]=28.12, *p*=0.003, *η*^2^ = 0.84) and in the data cleaned using ICA artifact rejection procedures (F[1,5]=8.62, *p*=0.03, *η*^2^ = 0.63). The same interaction was present in the EEG signals in the theta band, in both the raw (F[1,5]=7.34, *p*=0.04, *η*^2^ = 0.59) and cleaned data (F[1,5]= 8.57, p=0.03, *η*^2^ = 0.63). While a similar pattern was observed in the alpha band signals, there were no significant interactions for alpha, beta or gamma (all *p*>0.18).

**Figure 2.**
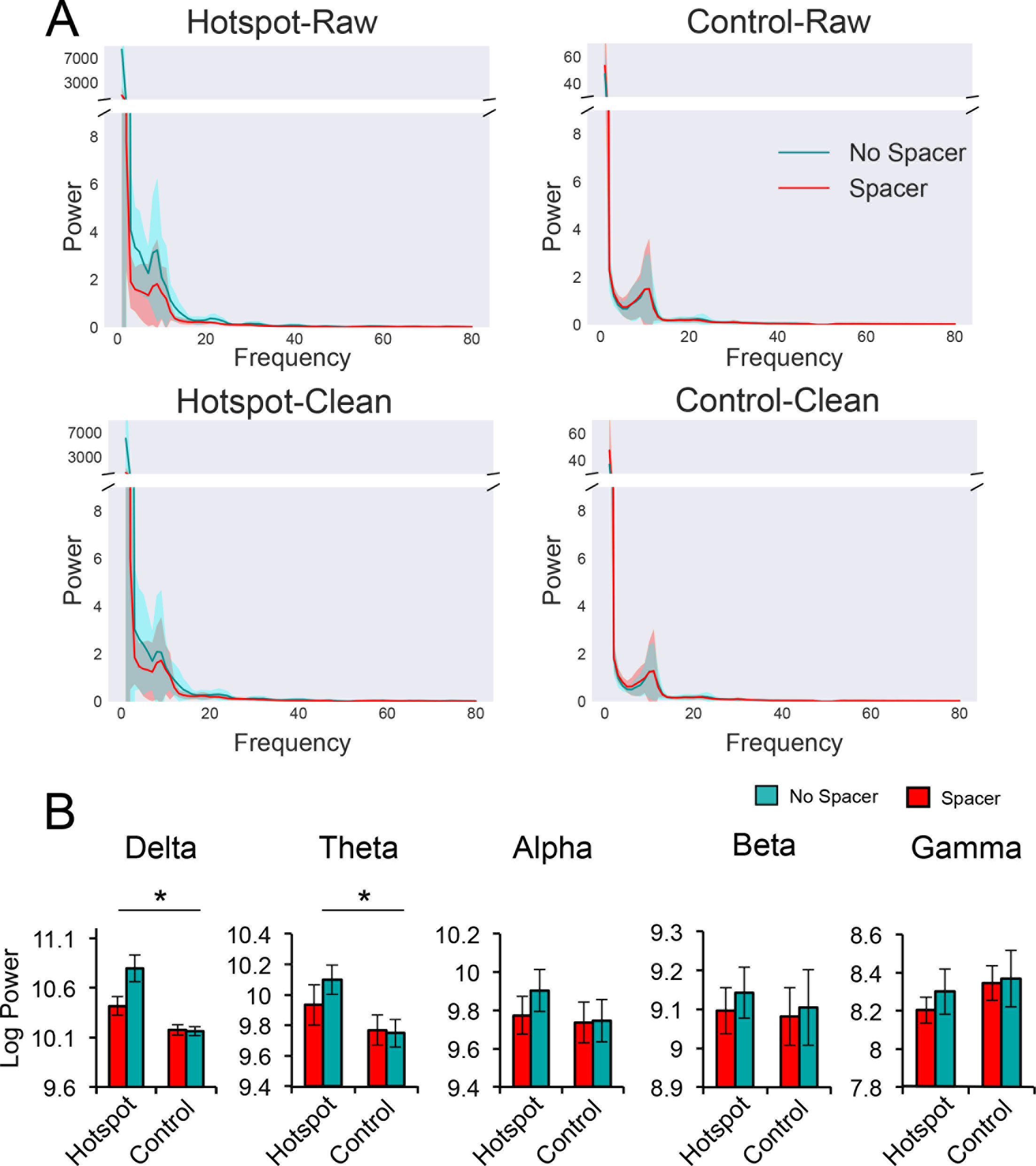
Power spectrum differences with and without spacer. Panel A depicts group average power spectra over 20 epochs for the hotspot electrode over which the coil and spacer were placed, and a control electrode (corresponding position on the opposite hemisphere). Signals are shown both raw (top panels) and cleaned using ICA (bottom panels). Shaded regions indicate standard deviation. Large artifacts manifest as greatly increased power in the ‘no spacer’ spectra (cyan) in the low frequency range when the TMS coil is in contact with the cap. This is the case in both the raw and cleaned data. With the inclusion of the spacer, artifacts are minimized and signals are more similar to those recorded from the control electrode. Panel B shows group level results of 2x2 factorial design ANOVA models, separated into five frequency bands. Bars depict mean logged power values. Error bars show standard error of the mean. Lines with a * indicate a significant ‘electrode’ x ‘spacer/no spacer’ interaction.

Additionally we tested whether some portion of the contact artifact may be attributed to the conduction of electrical noise through the TMS coil into the EEG electrodes. Supplementary Figure 1 (Panel A) shows the spectrums presented separately for the Spacer and No Spacer blocks, where it can be seen that there is no difference between the conditions where the TMS unit was switched ON and OFF. Panel B shows the same data presented instead contrasting the Spacer and No Spacer conditions, where it can be seen that the amplitude of signals in the low frequency range is higher when No Spacer was used (direct contact of TMS coil and EEG cap) in both situations (TMS machine ON and OFF). The fact that the artifact observed in the low frequency range occurs in the EEG signal even when the TMS unit is completely switched off, indicates that it is due to contact of the TMS coil on the cap and not from electrical noise conducted through the TMS apparatus.

## DISCUSSION

Here we demonstrate that artifacts arising in the pre-TMS EEG period from contact of the TMS coil on the surface of the cap are substantial, and exhibit frequency profiles that are well within the physiological range of viable brain signals. When the TMS coil was in direct contact with the EEG cap, the amplitude of EEG signals was elevated across all frequency bands, but evidently the lowest recorded frequencies were most susceptible, with those in the Delta and Theta range seriously affected. This can be seen from the very large effect sizes for the interactions between conditions with and without the spacer present and electrode position for these frequency bands. Importantly, standard data processing using ICA cleanup was insufficient to remove these artifacts, most likely because they closely resemble true electrophysiological data. The introduction of a plastic ‘coil spacer’ reduced this signal disturbance, and resulted in data more closely resembling that recorded from the corresponding (control) electrode in the opposite hemisphere.

Even without physical contact with the scalp, the fast changing magnetic field held near the hotspot can induce a flow of current in the underlying neural tissue. While many recent TMS-EEG investigations have proceeded with the TMS coil placed directly on the head (which is infact the configuration advertised commercially in several brochures for TMS-compatible EEG systems), the majority of well controlled investigations have implemented holding apparatus for positioning the coil over the head, advocating a ‘no contact’ approach (Ilmoniemi and Kičić 2009; Veniero et al. 2009), whereby the coil hovers close to the scalp, or introducing a foam layer between the coil and EEG cap (eg. Massimini et al. 2005). While coil hovering is one possible solution to the movement artifact problem, it is difficult to maintain over long testing sessions as natural subject movement causes the gap between the coil and the head to be non uniform over time, and contact can often occur. The spacer can be used with or without an additional coil holding apparatus, and ensures that the distance between scalp and coil remains fixed, even during subject head movements. While not specifically tested in this investigation, the Spacer may also contribute to a reduction in the bone conduction of the ‘click’ produced by the coil, and also reduce the mechanical forces produced by the vibration after the magnetic pulse, that propagate into the scalp.

### Source of the contact artifact

We have shown that electrical noise conducted through the TMS coil into the EEG system does not play a major role in the contact artifact, as it is present even in conditions where the TMS unit is switched off. However, movement factors such as human sway and limb or head positional drift, may produce low frequency artifacts. Additionally, transient hand contact with the electrode and friction during slippage of the coil may present as higher frequency noise, which we also observed to a lesser extent. It is also possible that the artifact is partly due to better conductance of the EEG signal as the pressure of the coil brings it closer to the scalp. Thus, we use the term ‘contact artifact’ to encompass the several different sources that are likely to contribute additively to the observed EEG signal disturbance when the coil is placed directly on the EEG cap.

### Limitations & Future directions

While the current version of the spacer has been adapted for use with the EGI cap system, small modifications may be required for use with other EEG systems. In particular, caps that have fully closed surfaces (rather than the open net-like design of EGI) may encounter more difficulties, as it is unknown whether contact of the spacer legs on the cap surface would transfer some portion of the movement artifacts to the nearby electrodes. However, with any existing EEG system it is expected that using the spacer to raise the coil a small distance above the electrodes and eliminating direct contact would result in higher quality data. We provide a fully editable 3D printer design file to allow other groups to make changes where necessary to accommodate their specific cap layout. It may be that the spacing of the tripod feet could be increased to reduce tension placed by the spacer on tight knit cap designs. Also, further modifications can be made post-printing, as perhaps it may be beneficial for certain types of cap to add a layer of rubber tape to the bottom of the tripod feet in order to reduce slippage on smoother cap surfaces, or to the surface of the TMS coil to avoid slippage against the spacer platform. Another limitation is that with the inclusion of the spacer, the TMS intensity required to evoke motor responses in the finger muscles was increased by 11% on average compared to when the coil was directly on top of the EEG cap. In some cases, the necessity to use higher intensities may prevent the participation of subjects with high RMTs, as the coil is more likely to overheat at high intensities. This is an unavoidable consequence of the extra distance between the scalp and the coil, and is a problem that is also present when using a coil holding apparatus with a similar distance air-gap between the coil and the head.

An additional point to note concerning the current investigation is the use of the EGI brand EEG system, which is designed to be ‘high impedance’ and to record good quality EEG data with higher than normal impedances (up to 50 kΩ). It is known that movement artifacts are amplified at high impedance (Ilmoniemi et al. 2009), and as such it may be the case that the contact artifact is less extreme with low impedance systems as what is portrayed here.

A challenge for this type of investigation using an in-vivo measure is the superposition of physiological signals with the artifactual signals. While we have endeavoured to isolate the dynamics of the contact artifact in conscious humans, and aimed to demonstrate that the Spacer is applicable in a real-laboratory context, a cleaner approach to characterise the artifact may involve repeating the measurements on a model head or realistic phantom. This would remove the complication of fluctuating physiological signals and isolate purely the artifact associated with the TMS coil contact.

### Conclusions

We introduce the ‘coil spacer’ for use in future simultaneous TMS-EEG recordings, to provide quality data from the hotspot region that is unaffected by movement artifact arising from contact between the coil and electrodes. We profile the extent of the low frequency movement artifacts that arise when no precautions are taken to avoid contact, and demonstrate the efficacy of a simple solution to the problem.

## Acknowledgements

This work was supported by Swiss National Science Foundation 320030_149561 and 320030_146531. We would also like to thank Andres Nussbaumer for help during data collection.

